# Reappearing Sensory Input Guides Visual Working Memory Prioritization

**DOI:** 10.1101/2024.02.07.579262

**Authors:** Damian Koevoet, Christoph Strauch, Marnix Naber, Stefan Van der Stigchel

## Abstract

Adaptive behavior necessitates the prioritization of the most relevant information in the environment (external) and in memory (internal). Internal prioritization is known to guide the selection of external sensory input, but the reverse may also be possible: Does the environment guide the prioritization of memorized material? Here we addressed whether reappearing sensory input can facilitate the prioritization of other non-reappearing memorized items held in visual working memory (VWM). Participants (total *n* = 96) memorized three orientations. Crucially some, but not all, items maintained in VWM were made available again in the environment. These reappearing items never had to be reproduced later. Experiment 1 showed that the reappearance of all but one memory item benefited accuracy and speed to the same extent as a spatial retro cue. This shows that reappearing items allow for the dynamic prioritization of another non-reappearing memorized item. What aspects of the reappearing sensory input drive this effect? Experiments 2-4 demonstrated that prioritization was facilitated most if reappearing items matched VWM content in terms of both location and orientation. Sensory input fully matching VWM is possibly processed more efficiently and/or protects against interference, ultimately leading to stronger prioritization of other memory content. We propose that the link between sensory processing and VWM is bidirectional: internal representations guide the processing of sensory input, which in turn facilitates the prioritization of other VWM content to subserve adaptive behavior. All data and analysis scripts are available here: https://osf.io/qzvkc/.

## Introduction

The world provides us with overwhelming visual input that cannot all be processed simultaneously. Visual attention allows for the selective processing of the most relevant visual input (Carrasco, 2011; Posner, 1980). Attention cannot only select external information, but memoranda held in visual working memory (VWM) can be prioritized by attention as well. Following van Ede and Nobre (2023, p. 139), we here define VWM prioritization as “the ability to change the priority or accessibility among multiple coexisting internal representations in working memory”. As such, VWM prioritization has been shown to improve behavioral outcomes such as accuracy and/or response speed (Griffin & Nobre, 2003; Heuer et al., 2020; Koevoet, Strauch, et al., 2024; Landman et al., 2003; Olivers et al., 2011; Olivers & Roelfsema, 2020; Souza & Oberauer, 2016; van Ede, 2020; van Ede & Nobre, 2023). Although overwhelming sensory input poses a challenge, we can leverage this input to guide us to attend the most important external information (e.g. a rapidly approaching car, see Võ et al., 2019). Here, we addressed whether sensory input may also be used to guide the prioritization of VWM content.

How could sensory input facilitate the prioritization of memorized items? Imagine you are riding your bicycle in a busy street and you want to cross the road (Figure 1). You first look to your left, and see a black, a red and a black car approaching. These vehicles are important, and you store them into VWM for later reference. You then take a look to the right where no vehicles approach, but while doing so the black and red car approaching from the left pass your view. The reappearance of these cars signals that now solely the black car remains relevant to cross the road safely. What happens to the memory representation of the black car? Although only cars reappeared that are now obsolete, these cars may guide one to internally prioritize the black car. This leads to the – initially counter-intuitive – proposition that you remember the remaining car better, even though it did not reappear. Here, we investigated whether reappearing sensory input can indeed facilitate the prioritization of VWM content.

The idea that availability in the external world affects VWM use is well established. If relevant information is available in the environment, participants tend to not load up VWM fully (e.g. Ballard et al., 1995; Böing et al., 2023; Draschkow et al., 2021; Hoogerbrugge et al., 2023; Kvitelashvili & Kessler, 2024; Somai et al., 2020) – or sometimes not at all (Chota et al., 2023). Instead, participants often prefer to resample to-be-memorized material and only encode one or two items at a time (Aivar et al., 2005; Ballard et al., 1995; Böing et al., 2023; Chota et al., 2023; Draschkow et al., 2021; Droll & Hayhoe, 2007; Grinschgl et al., 2021; Hoogerbrugge et al., 2024; Hoogerbrugge et al., 2023; Koevoet, Naber, et al., 2023; Kvitelashvili & Kessler, 2024; Melnik et al., 2018; Meyerhoff et al., 2021; O’Regan, 1992; Risko & Gilbert, 2016; Sahakian et al., 2023a, 2023b; Somai et al., 2020; Van der Stigchel, 2020; L. Xu et al., 2023). Put differently, participants typically prefer to simply look at externally available items over storing them in VWM, likely because this is less effortful (Risko & Gilbert, 2016; Van der Stigchel, 2020) (although sampling also has a cost, see Koevoet, Strauch, et al., 2023; Koevoet, Van Zantwijk, et al., 2024). In contrast, whenever external information becomes less accessible (i.e. further away or longer delay times before access), participants shift toward storing more in VWM (e.g. Draschkow et al., 2021; Hoogerbrugge et al., 2023; Sahakian et al., 2023a). These observations demonstrate that VWM use is flexibly adjusted to the constraints set by the environment.

While previous studies demonstrate a robust link between external availability and VWM encoding, the to-be-memorized material was highly stable in most tasks (Draschkow et al., 2021; Grinschgl et al., 2021; Hoogerbrugge et al., 2023; Koevoet, Naber, et al., 2023; Melnik et al., 2018; Meyerhoff et al., 2021; Sahakian et al., 2023a, 2023b; Somai et al., 2020) (but see Aivar et al., 2005; Hoogerbrugge et al., 2024; L. Xu et al., 2023). This contrasts many situations in everyday life in which the world is highly dynamic as items can appear, disappear and reappear from the visual field within a matter of seconds (Nobre & van Ede, 2023). Humans may be able to capitalize on the dynamic nature of the world by using it to prioritize relevant memorized material. Through reappearance, sensory input may facilitate operations of VWM besides encoding, such as prioritization. Here, we report four VWM experiments that address this possibility. We mimicked a dynamic scenario by making some but not all maintained items reappear, which may facilitate VWM prioritization of other externally unavailable content to guide goal-directed behavior (Figure 1) - essentially acting as a retro-cue. We hypothesized that the reappearance of maintained items that were then obsolete, would allow for the prioritization of remaining VWM content.

**Figure 1:**
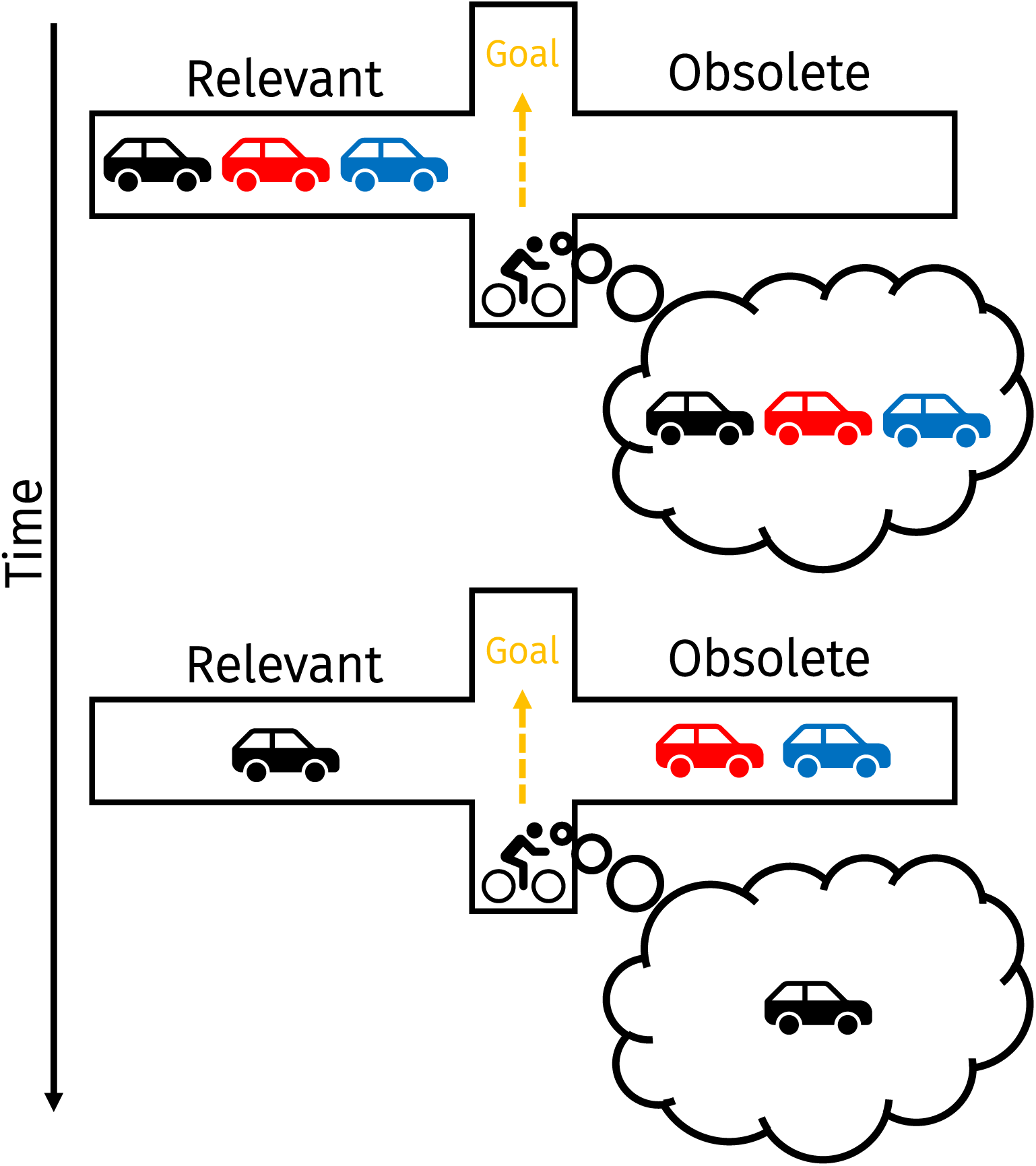
An example of how sensory input conveys information that might facilitate internal prioritization. In a dynamic environment, relevant information can become obsolete in a matter of seconds. When trying to cross the road, three cars are approaching from the left. Upon looking to the right where no cars are approaching, the black and red cars pass by. The black and red cars have now become obsolete and only the black car remains relevant to your goal of crossing the road. We here studied whether such memory-matching sensory input can be used to boost remaining VWM content through prioritization.

## Experiment 1

### Methods

#### Transparency and openness

All data and analysis scripts are available on the Open Science Framework: https://osf.io/qzvkc/. Data for Experiments 1-3 were collected in 2023, and Experiment 4 was conducted in 2024. None of the experiments were preregistered.

#### Participants

Twenty-four participants (including the first author) with normal or corrected-to-normal vision took part in Experiment 1 (*M_age_* = 24.33, range: 19-29; 8 males; 3 left-handed). We used G*Power (v3.1) to conduct a power analysis (Faul et al., 2007). As we did not have an adequate estimate of an effect size from the previous literature for our effect of interest, we chose a medium effect size of 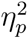 = .06. To detect a main effect of this size with .80 power at an *α* = .05 with four conditions (within-subjects), 23 participants were required. We tested one additional participant for adequate counterbalancing (see below). The final sample size is comparable with, or even exceeds, previous work investigating VWM prioritization (e.g. Gunseli et al., 2015; van Ede et al., 2020), and studies demonstrating the relationship between external availability and VWM use (e.g. Hoogerbrugge et al., 2024; Hoogerbrugge et al., 2023; Koevoet, Naber, et al., 2023; Somai et al., 2020). Participants reported no history of attention-deficit related disorders or autism spectrum disorder. All participants were compensated with €8 or course credits. The first author (D.K.) participated in Experiments 1-3. To assess whether this affected our conclusions, we conducted all analyses from these Experiments while also excluding the first author’s data. Note that all outcomes and conclusions remain unaltered when excluding this participant unless otherwise explicitly stated (which occurred for only a single comparison). The experimental procedure was approved by the local ethical review board (approval code: 21-0297).

#### Stimuli

Memory stimuli were three Gabor gratings (2.5°x2.5° in width x height; spatial frequency: 3 cpd) with a random orientation, positioned at three constant locations in a triangle at an eccentricity of 4° from the center of the screen (see Figure 2). Mask stimuli were concentric gratings (2.5°x2.5°; 3 cpd) and were always presented at the same locations as the Gabor gratings. Arrow stimuli (0.5°x1.5°) consisted of a straight black line with two ∧ shapes either pointing toward the same direction (spatial retro cue block) or toward each other (no spatial cue; all other blocks). The response wheel (4°x4°) consisted of one larger circle on which two smaller circles were positioned. Participants rotated the orientation of these smaller circles to reproduce the memorized orientation of the stimuli using the mouse (as in Gresch et al., 2024; van Ede et al., 2020; van Ede et al., 2019). The black outline of an otherwise transparent square (2.5°x2.5°) probed the location of the item which orientation should be reported. Stimuli were presented on a gray background (RGB value: [128, 128, 128]) with a DELL U2417H monitor (1920 x 1080; 60 Hz) using PsychoPy (v2021.2.3).

**Figure 2:**
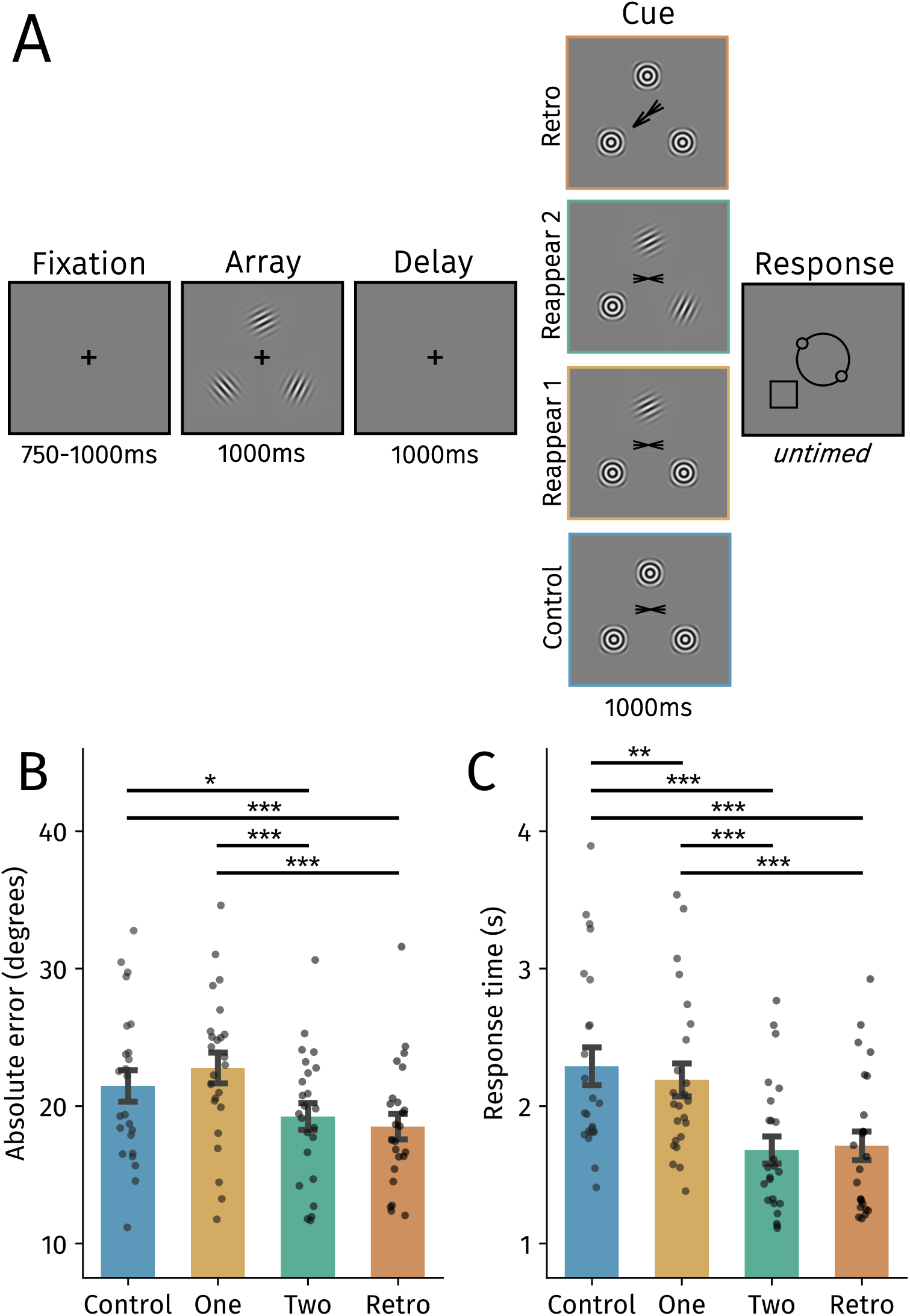
**a**)Trial structure of Experiment 1. After a fixation period, participants encoded the orientation of three Gabor gratings. Different cues were presented after a delay period depending on the block. In the reappear-two block, participants could infer which item should be reported because the reappearing stimuli would never be probed at the end of the trial. The reappear-one block left participants with two remaining options of possible orientations to report. Using an arrow cue, participants shifted attention toward the to-be-probed item in the retro cue block. Lastly, participants did not receive any information during the cue phase in the control block. **b)** Mean absolute error in degrees across block types. **c)** Average response times across block types. Black points reflect individual participants. Error bars represent SEM. Holm-corrected: \**p* < .05, \*\**p* < .01, - *p* < .001.

#### Procedure

Participants were positioned approximately 60 cm from the screen in a chin- and head-rest. Trials started with a brief fixation period (750-1000 ms). Participants encoded the orientations (all random) of three Gabor gratings that were presented for 1000 ms. After the delay period (1000 ms), a cue was presented (1000 ms) that differed between blocks. In the reappear-two block, two memorized gratings reappeared in their original location which indicated that these orientations would not be probed at the end of the trial (100% valid). The reappear-one block was identical to the reappear-two block, but only one grating reappeared. The retro cue block provided participants with an arrow pointing toward the to-be-reproduced orientation. The control block was identical to the arrow cue block but there was no informative cue. To keep visual input similar across blocks, masks were always presented on locations where no gratings reappeared and a non-informative cue was shown at fixation (except in the retro cue block). After the cue phase, participants reproduced the orientation that was spatially probed (indicated by the black square) as precisely as possible by moving the mouse to turn the response wheel, and confirmed with a left mouse click. The starting position of the mouse cursor (3° eccentricity) and thus the orientation of the response wheel were randomized on every trial. Blank inter-trial intervals varied randomly between 500-1000 ms.

Participants completed one block of trials per condition. Each block consisted of 60 trials and the order of blocks was counterbalanced using a balanced Latin square design across participants. To reduce fatigue, participants took a break after every block. Participants completed five practice trials of every block before the experiment.

#### Data Analysis

All processing and analyses were performed using custom Python (v3.9.7) scripts. Practice trials and trials with responses faster than 200 ms or slower than 6000 ms were not considered for analysis (1.1% of trials). A total of 5,697 trials were entered into the analyses.

The absolute error of the orientation report was used as a measure of accuracy (as in e.g. Gresch et al., 2024; Koevoet, Naber, et al., 2023; van Ede et al., 2020; van Ede et al., 2019). Although ultimately no participants were excluded from Experiment 1, we took two steps to ensure participants could perform the task above chance level. First, participants were excluded if the average absolute error was equal or higher than 45° in any of the conditions (as in Gresch et al., 2024). Second, across all conditions we shuffled participants’ responses and calculated the averaged shuffled absolute error. We compared this shuffled absolute error to participants’ actual average error 10,000 times and determined participants were better than chance if their actual responses led to lower absolute errors than the shuffled responses in at least 95% of the comparisons. No participants were excluded based on these criteria in Experiment 1, indicating that they were able to perform the task above chance level (all *p* < .001). Additionally, we also analyzed response times, which here represent the time from the response probe until the response was completed fully (i.e. the mouse click confirming the response). Note that both response accuracy and/or speed may be affected by VWM prioritization (van Ede & Nobre, 2023).

Accuracy and response time data were analyzed using linear mixed-effects models. We chose to use linear mixed-effects models over traditional statistical approaches because these models are more appropriate given the data structure, are more flexibly and have increased statistical power (Brysbaert & Stevens, 2018). Models were fit using the *statsmodels* package in Python.^1^ We modeled by-participant intercepts and slopes of cue type effects in every model to minimize Type 1 errors (Barr, 2013) - Wilkinson notation: Outcome ∼ Cue type + (1 + Cue type|Participant). All *p*-values from the linear mixed-effects models were corrected for multiple comparisons using the Holm method, and we report 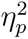 as a measure of effect size (as in Kumle et al., 2024).

## Results

To determine if participants used reappearing items to flexibly prioritize the most relevant VWM content, we analyzed absolute errors and response times (Figure 2). If reappearing gratings guide the prioritization of the remaining material, performance should be enhanced compared with the control block.

Indeed, participants were more accurate in the reappear-two block than in the reappear-one (*β* = 3.56 ± .75, *t* = 4.73, *p* < .001, 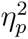 = .40) and control blocks (*β* = 2.21 ± .77, *t* = 2.86, *p* = .013, 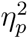 = .20) (Figure 2B). Similarly and in line with the literature (Myers et al., 2018; Souza & Oberauer, 2016; van Ede & Nobre, 2023), participants were more accurate in the retro cue condition than in the reappear-one (*β* = 4.30 ± .77, *t* = 5.58, *p* < .001, 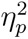 = .49) and control blocks (*β* = 2.95 ± .80, *t* = 3.68, *p* < .001, 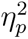 = .29), but absolute errors did not significantly differ between the reappear-two and retro cue blocks (*t* = .99, *p* = .323, 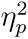 = .03). This indicates that two reappearing items guide VWM prioritization in a comparable manner to traditional retro cues. In contrast, absolute errors in the reappear-one condition did not differ significantly with the control condition (*t* = 1.79, *p* = .147, 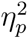 = .09).

The response time analysis tells a similar story (Figure 2C): participants responded faster in the reappear-two and retro cue blocks than in the reappear-one (reappear-two: *β* = 510.03 ± 29.50, *t* = 17.29, *p* < .001, 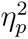 = .93; retro: *β* = 479.63 ± 41.94, *t* = 11.44, *p* < .001, 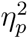 = .85) and control blocks (reappear-two: *β* = 607.27 ± 41.95, *t* = 14.48, *p* < .001, 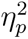 = .90, retro: *β* = 576.86 ± 56.93, *t* = 10.13, *p* < .001, 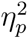 = .82). The reappear-two and retro cue blocks did not differ significantly (*t* = 1.03, *p* = .30, 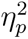 = .04), and participants responded faster in the reappear-one than in the control block (*β* = 97.23 ± 29.58, *t* = 3.29, *p* = .002, 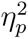 = .32). For the comparison in accuracy between the reappear-two and retro cue blocks, we also conducted a Bayesian linear mixed-effects model to potentially obtain evidence for the null hypothesis. To this end, we fit models (using the *brms* package in *R*) and computed a Bayes factor for the comparison of interest. We obtained evidence for the null-hypothesis (BF_01_ = 3.23), indicating that accuracy was similar between the two conditions.

Together, these analyses show that participants were able to use the reappearing gratings to prioritize the most relevant VWM content. More specifically, when two gratings reappeared, participants responded more accurately and faster than in the control condition. Behavior in the reappear-two condition was comparable to that in a traditional spatial retro cue. However, whenever one grating reappeared, accuracy remained somewhat comparable (but decreased numerically) although responses were faster compared with the control condition. In terms of our aforementioned example (Figure 1), this implies that humans can prioritize the black car whenever the black and red cars have passed. If instead only the black car has passed, behavior is not guided as effectively by internally focusing on both the black and red car - as responses were slightly faster but not more accurate. These results resonate somewhat with the idea that only a single item held in VWM can guide behavior at a time (Olivers et al., 2011; van Moorselaar et al., 2014) (but see Beck et al., 2012; Frătescu et al., 2019; Hollingworth & Beck, 2016; J. R. Williams et al., 2022, also see Discussion). It is also possible that the singleton grating in the reappear-1 condition captured attention (Theeuwes, 1994), which caused interference during the internal prioritization process. Regardless of what may explain the lack of an accuracy benefit in the reappear-1 condition, our findings demonstrate that dynamic sensory input can facilitate the prioritization of at least one VWM representation to guide behavior.

## Experiment 2

The data from Experiment 1 showed that participants were able to use reappearing items to prioritize non-reappearing content held in VWM. These reappearing items consisted of two main features: an orientation and a location. Many studies suggest that VWM predominantly employs spatial reference frames whenever possible - even when location is task-irrelevant (e.g. Draschkow et al., 2022; Foster et al., 2017; Luria et al., 2016; Treisman & Zhang, 2006; van Ede et al., 2019), which may suggest that participants predominantly used location information to guide prioritization in Experiment 1. Nonetheless, orientation information matching VWM content may also contribute to the prioritization process.

This latter possibility is supported by neural and psychophysiological evidence. For instance, sensory recruitment models posit that working memory and perception recruit the same low-level sensory brain areas (Chota & Van der Stigchel, 2021; Christophel et al., 2017; Harrison & Tong, 2009; Serences, 2016; Serences et al., 2009) (but see Y. Xu, 2020, 2023). Neuroimaging and pupillometric studies have supported this notion by revealing enhanced responses to visual stimuli matching the content of VWM, regardless of their location (Gayet et al., 2017; Olmos-Solis et al., 2018; Wilschut & Mathôt, 2022) (also see Karabay et al., 2024). Moreover, what is stored in VWM guides attention (Olivers et al., 2006; Soto et al., 2008), accelerates access to awareness (Ding, Naber, Paffen, Gayet, et al., 2021; Ding, Naber, Paffen, Sahakian, et al., 2021; Gayet et al., 2013), and affects eye-movements (Hollingworth et al., 2013; Silvis & Van der Stigchel, 2014). Gayet et al. (2017) suggested that VWM pre-activates neuronal populations responsible for visual perception, leading to enhanced processing of matching visual input. Through such memory-driven pre-activation, reappearing items matching memory in terms of both orientation and location may be processed deeper and/or faster than reappearing items not matching the memorized orientations. Enhanced sensory processing of the fully memory-matching reappearing items could make it easier to deduce which orientation should be reported, leading to more effective VWM prioritization. In Experiment 2, we investigated which aspects - orientation, location or a combination - of reappearing items facilitate VWM prioritization most effectively.

## Methods

The methods in Experiment 2 were identical to Experiment 1 unless specified explicitly.

### Participants

Twenty-seven participants with normal or corrected-to-normal vision took part in Experiment 2. Three participants were excluded for not performing above chance level (see Methods Experiment 1). This left a final sample of twenty-four participants (including the first author) as in Experiment 1 (*M_age_* = 23.00 (range: 18-30), 8 males, 6 left-handed) that were able to perform the task above chance level (all *p* < .001). We chose the same sample size and number of trials per condition as in Experiment 1 for comparability, and because we did not expect to find smaller effect sizes in Experiment 2.

### Procedure

Participants completed four blocks containing differing cues: combination, orientation, location and control (Figure 3). One difference from the first experiment is that the spatial probe (the black square) in the response screen was removed for all blocks besides the control block. We removed this probe in the cue blocks because 1) this ensured participants had to use the cues to deduce which orientation should be reported, 2) the spatial probe may induce reliance on location information, and 3) the probe would likely confuse participants in the orientation cue block due to inconsistent positions of the gratings during the encoding and cue phases.

**Figure 3:**
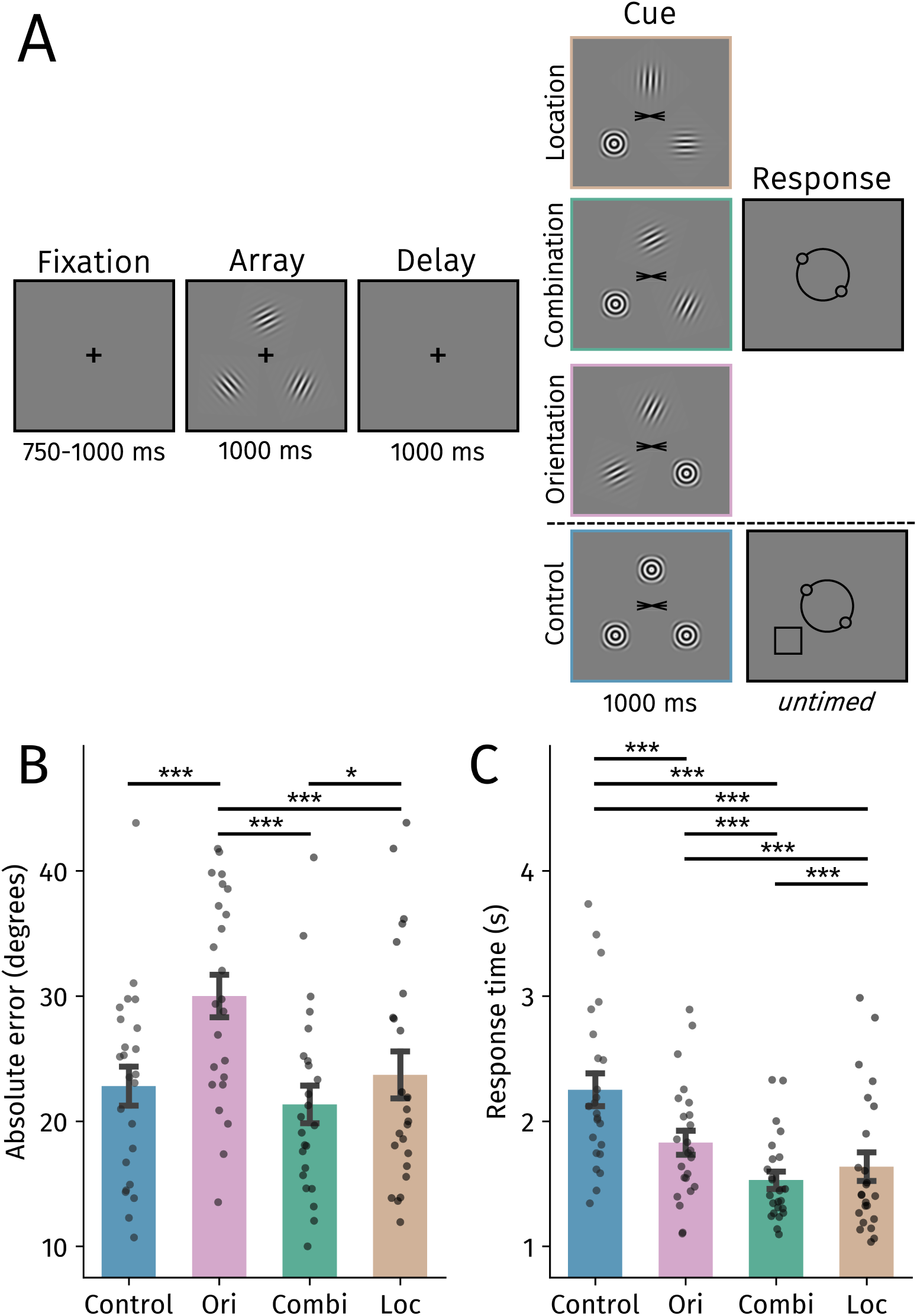
**a**) Trial structure of Experiment 2. In this example, the bottom left orientation (’\’) has to be reproduced. In the combination block, participants deduced which item should be reported using a combination of location and orientation information from the cue. In contrast, participants could only use the location or the orientation information in the other experimental blocks. Participants used a spatial probe during the response screen to infer what item should be reported in the control block.. **b)** Mean absolute error in degrees across block types. **c)** Average response times across block types. Black points reflect individual participants. Error bars represent SEM. *Abbreviations:* Ori = Orientation, Combi = Combination, Loc = Location. Holm-corrected: \**p* < .05, \*\**p* < .01, - *p* < .001.

The combination block was identical to the reappear-two block in Experiment 1, but there was no spatial probe. The location block was similar but although two stimuli reappeared in congruent locations, they now had a random orientation. This way participants could only use the location and not the orientation information to infer which item should be reported. In contrast, in the orientation block two stimuli reappeared with an identical orientation as before but now in a random location. Here, participants relied exclusively on the orientation information and not the location to deduce which orientation should ultimately be reported. The control block was identical to Experiment 1.

After discarding trials (0.5%) with very fast (<200 ms) or very slow (>6000 ms) response times, 5,790 trials were retained for the analyses.

## Results

If both the orientation and location information from the reappearing gratings facilitate prioritization, performance should be best in the combination block compared with the other blocks wherein items reappeared. Absolute errors were indeed smaller in the combination than in the location (*β* = 2.36 ± .86, *t* = 2.74, *p* = .019, 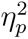 = .25) and orientation blocks (*β* = 8.66 ± .86, *t* = 10.03, *p* < .001, 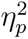 = .81) (Figure 3B). When participants relied exclusively on spatial information, they were more accurate than when only using orientation information (*β* = 6.30 ± .99, *t* = 6.39, *p* < .001, 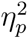 = .64). Moreover, reports were more precise in the control compared with the orientation block (*β* = 7.21 ± .87, *t* = 8.34, *p* < .001, 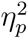 = .75). Accuracy in the combination and location blocks did not significantly differ from the control block (*t*s < 1.47, *p*s > .28, 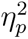s < .10).

The response time analyses complemented the accuracy findings and ruled out potential speed-accuracy trade-off effects for almost all comparisons (besides those with the control block). Participants responded fastest in the combination block (Figure 3C; location: *β* = 106.98 ± 30.24, *t* = 3.57, *p* < .001, 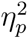 = .35; orientation: *β* = 300.70 ± 30.25, *t* = 9.94, *p* < .001, 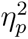 = .81; control: *β* = 720.70 ± 43.92, *t* = 16.41, *p* < .001, 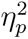 = .92). This is in line with the accuracy analysis, and again indicates that a combination of orientation and location information facilitates VWM prioritization most effectively. Additionally, responses were faster during the location block compared with the orientation block (*β* = 193.73 ± 43.91, *t* = 4.41, *p* < .001, 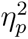 = .46), indicating that location guided prioritization more effectively than orientation information in the current task.^2^ Response times were slower in the control than in the orientation (*β* = 420.00 ± 30.28, *t* = 13.87, *p* < .001, 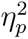 = .89) and location blocks (*β* = 613.73 ± 60.14, *t* = 10.21, *p* < .001, 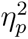 = .82).

Our results suggest that reappearing items matching VWM guide the prioritization of the remaining maintained grating most effectively. A combination of location and orientation information led to enhanced accuracy and speed when compared with the other blocks wherein gratings reappeared. Location was more effective than orientation information in facilitating VWM performance. In contrast to Experiment 1, we did not observe a difference in accuracy between the control and combination blocks. We suspected that this may be attributed to the spatial probe (i.e. black square) during the response screen in the control block but not in the other blocks. Participants may have used this probe as a spatial pointer to facilitate task performance when compared with the other blocks. Another possibility is that the inclusion of the orientation block – wherein items reappeared with the same orientation though not at their original locations - may have encouraged participants to encode orientations unbound to their spatial locations, and this encoding strategy may have extended to other blocks. We addressed this, and other remaining issues, in Experiment 3.

## Experiment 3

Experiment 3 aimed to address two issues. First, we assessed the robustness of the main findings from the first two experiments (while always including the spatial probe during recall). Second, we controlled for a potential effect of low-level visual adaptation by briefly presenting masks after the encoding phase. When not controlling for low-level visual adaptation, it may be possible that fully-matching reappearing gratings are easier to see, just because they were presented briefly before - not because they are held in memory. Therefore, we used masks to ameliorate potential effects of visual adaptation to ensure we are tapping into memory-related processes.

## Methods

The methods in Experiment 3 were identical to Experiment 2 unless specified explicitly.

### Participants

In Experiment 3, twenty-five participants with normal or corrected-to-normal vision took part. One participant was excluded for not performing above chance level (see Methods Experiment 1). This left a final sample of twenty-four participants (including the first author) as in Experiment 1 and 2 (*M_age_* = 22.08 (range: 19-27), 3 males, 1 left-handed) that were able to perform the task above chance level (all *p* < .001). As we aimed to replicate the main findings from Experiments 1 and 2, we chose the same sample size and number of trials per condition as in those experiments.

### Procedure

The experiment consisted of three blocks (Figure 4): the control, location and combination blocks from Experiment 2. To control for adaptation effects, masks were presented for 200 ms after memory array offset. Furthermore, the black square probe was included in all blocks to enhance comparability between them.

**Figure 4:**
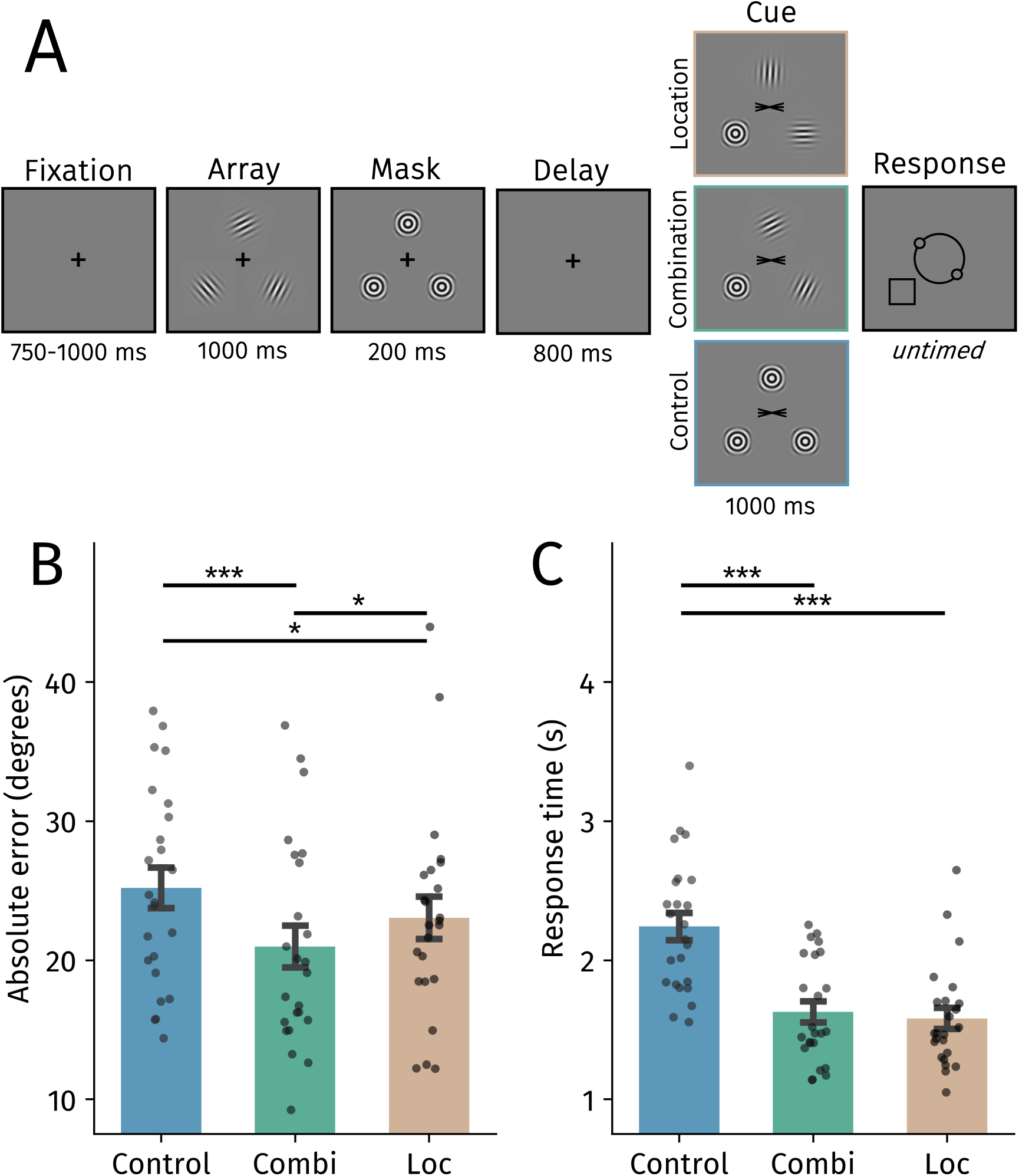
**a**) Trial structure of Experiment 3. **b)** Mean absolute error in degrees across block types. **c)** Average response times across block types. Black points reflect individual participants. Error bars represent SEM. *Abbreviations:* Combi = Combination, Loc = Location. Holm-corrected: \**p* < .05, - *p* < .001.

Trials (0.34%) with very fast (<200 ms) or very slow (>6000 ms) response times were discarded. 4,305 trials were retained for the analyses.

## Results

In Experiment 1 we observed that whenever two memory-matching gratings reappeared, participants were more accurate and faster when reproducing the cued orientation. We replicate both of these findings (Figure 4): participants were more precise (*β* = 4.23 ± .92, *t* = 4.60, *p* < .001, 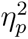 = .48) and responded faster (*β* = 612.69 ± 48.84, *t* = 12.55, *p* < .001, 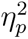 = .87) in the combination compared with the control block.

Experiment 2 demonstrated that especially gratings matching memory in terms of both location and orientation facilitated VWM prioritization. In line with this, responses were more accurate in the combination compared with the location condition (*β* = 2.07 ± .82, *t* = 2.53, *p* = .023, 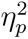 = .22). Location information did, however, also facilitate VWM prioritization as reports were slightly more precise (*β* = 2.16 ± 1.07, *t* = 2.03, *p* = .043, 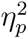 = .15)^3^ and faster in the location block than in the control block (*β* = 659.52 ± 68.52, *t* = 9.63, *p* < .001, 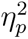 = .80). Response times did not significantly differ between the location and combination blocks (*t* = 1.48, *p* = .138, 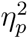 = .09). Together, Experiment 3 replicated the main findings from the previous experiments and ruled out low-level visual adaptation as a possible confound.

One discrepancy between Experiments 2 and 3 is that we found a significant difference in terms of accuracy between the location and control condition in Experiment 3 but not in Experiment 2. This could have been caused by the appearance of the black probe square in Experiment 3 but not 2 or possibly due to participants encoding the gratings less bound to space in Experiment 2 due to the inclusion of the orientation block (also see Experiment 2). However, when removing the data from the first author, this difference was non-significant. We conclude that this effect is likely not reliable (or small) and should be interpreted with caution.

## Experiment 4

Experiment 4 aimed to address two further questions. First, it is possible that participants prioritized the singleton mask stimulus without considering the reappearing gratings. Although unlikely, as we found consistent differences between location and combination conditions, we wanted to rule out this possibility experimentally. We did so by increasing the total number of locations and removed the mask stimulus on the to-be-probed location. This ensured that participants could not selectively prioritize the mask stimulus, but instead had to use the reappearing stimuli to infer the to-be-reported grating. Second, we included a condition where two mask stimuli appeared on locations that were never probed. This condition could not cause feature-based interference because no orientations were presented during the cue period. This allowed us to study the potential role of interference caused by ‘new’ orientation information that was presented when gratings reappeared in the location but not the combination condition Note that the masks condition provides similar information as a spatial retro cue.

## Methods

The methods in Experiment 4 were identical to Experiment 3 unless specified explicitly.

### Participants

Twenty-eight participants with normal or corrected-to-normal vision took part in Experiment 4. Four participants were excluded for not performing above chance level (see Methods Experiment 1). This left a final sample of twenty-four participants as in Experiments 1-3 (*M_age_* = 22.13 (range: 19-28), 4 males, 1 left-handed) that were able to perform the task above chance level (all *p* < .001).

### Procedure

The experiment consisted of three blocks (Figure 5A): slightly adapted versions of the location and combination blocks from Experiment 2 and 3, and a masks condition. In the masks condition, two mask stimuli without orientation information reappeared, effectively acting like a spatial retro cue. As in the previous experiments, stimuli that originally occupied these locations were never probed at the end of the trial. This allowed us to test whether or not the memory-match effect affected by the interference caused by ‘new’ gratings in the location condition. Another important task design difference is that stimuli could now be presented in six, instead of three different locations compared with the previous experiments. Moreover, no stimulus was now presented on the to-be-probed location. This ensured that participants could not selectively prioritize the mask stimulus without considering the reappearing gratings.

**Figure 5:**
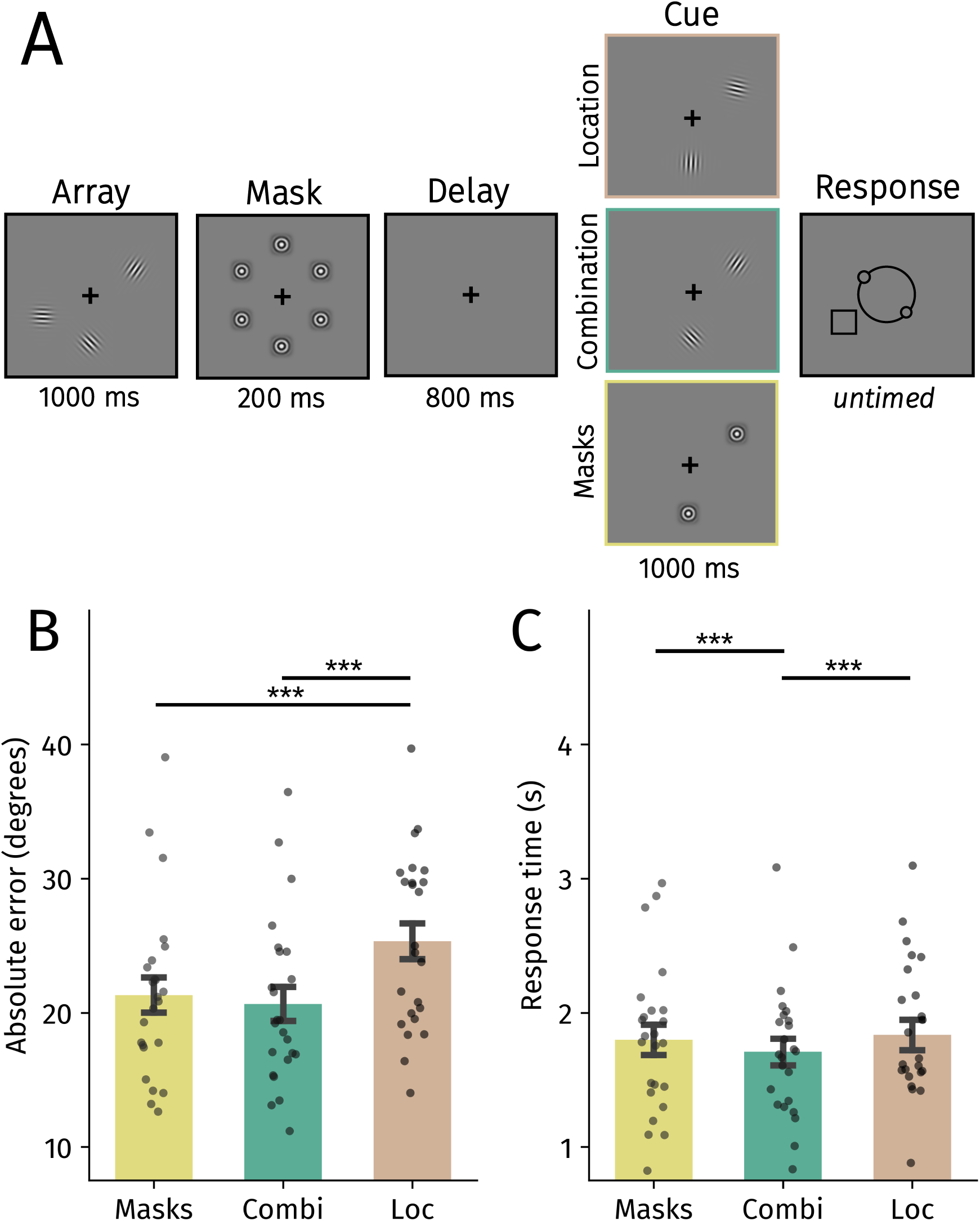
**a**) Trial structure of Experiment 4. **b)** Mean absolute error in degrees across block types. **c)** Average response times across block types. Black points reflect individual participants. Error bars represent SEM. *Abbreviations:* Combi = Combination, Loc = Location. Holm-corrected: - *p* < .001.

We increased the number of trials per condition to 75, slightly increasing statistical power. As in the previous experiments, we discarded trials (0.76%) with very fast (<200 ms) or very slow (>6000 ms) response times. The analysis included 5,359 trials in total. Note that for the response time analysis, the model did not converge when including by-participant random slopes, and we therefore fit the model as follows, Wilkinson Notation: Response time ∼ Cue Type + (1|Participant) (as recommended by Barr, 2013).

## Results

In Experiment 2 and 3, we observed that participants were more accurate in the combination than in the location condition. Using an improved design that addressed possible caveats, we replicated the same pattern (*β* = 4.69 ± .77, *t* = 6.11, *p* < .001, 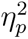 = .62) (Figure 5B). Ruling out a speed-accuracy trade-off, participants responded faster in the combination than in the location condition (*β* = 127.44 ± 23.11, *t* = 5.51, *p* < .001, 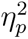 = .57) (Figure 5C).

We then turned to the masks condition. The masks cue provided the same spatial information as the other conditions, but did not match the content held in memory and could not cause interference by introducing new orientation information. This allowed us to study the potential effect of presenting new orientation information (Figure 5). Participants were more accurate in the masks compared with the location condition (*β* = 4.03 ± 1.13, *t* = 3.58, *p* < .001, 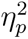 = .36), and response accuracies did not differ significantly between the masks and combination conditions (*β* = .66 ± .92, *t* = .72, *p* = .47, 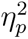 = .02).

As for response times, we found no significant difference between the masks and location conditions (*β* = 36.51 ± 23.13, *t* = 1.58, *p* = .11, 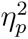 = .10). By contrast, participants responded faster in the combination than in the masks condition (*β* = 90.93 ± 23.11, *t* = 3.94, *p* < .001, 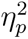 = .40).

Together, these results indicate that the accuracy difference between the location and combination conditions were, at least in part, driven by interference caused by new orientation information. Indeed, memory-matching reappearing gratings seem to not cause orientation-based interference. Accuracy performance was similar between the masks and combination conditions. This is in line with the finding from Experiment 1 where accuracy was similar in the reappear-2 and retro cue conditions. Note that reappearing gratings that matched VWM did yield faster response times compared with the mask stimuli. These results indicate that memory-matching reappearing gratings facilitate the prioritization of a non-reappearing item (at least) as effectively as a spatial retro cue.

## Discussion

We report four experiments that investigated whether reappearing sensory input facilitated the prioritization of another non-reappearing memorized item. Experiment 1 showed that reappearing memorized items facilitated the prioritization of a single remaining relevant item held in VWM. The beneficial effects on performance elicited by the reappearing items were comparable to a spatial retro cue. In Experiment 2, 3 and 4, we went a step further and asked what aspects of reappearing items guide VWM prioritization most effectively. We observed that items matching VWM in both orientation and location guided internal prioritization most strongly (compared with only the orientation or location). Experiment 4 demonstrated that this fully-matching benefit observed in previous experiments may be attributed to the presentation of items with novel orientations. This indicates that memory-matching reappearing material may not be ‘special’ compared with a spatial retro cue, yet they may constitute a more naturally occurring instance of a way to guide internal prioritization. Our results demonstrate that humans are able to use reappearing sensory input to guide the prioritization of the most relevant VWM content.

VWM is typically studied by briefly presenting to-be-encoded material, and subsequently removing this information from the environment (e.g. Luck & Vogel, 1997; Vogel et al., 2006). Another line of work instead leaves to-be-memorized items externally available indefinitely to resemble more stable environments (Ballard et al., 1995; Böing et al., 2023; Chota et al., 2023; Draschkow et al., 2021; Grinschgl et al., 2021; Hoogerbrugge et al., 2023; Koevoet, Naber, et al., 2023; Meyerhoff et al., 2021; O’Regan, 1992; Risko & Gilbert, 2016; Sahakian et al., 2023a, 2023b; Somai et al., 2020; Van der Stigchel, 2020). Here we combined these approaches by using sensory input which appeared, disappeared and reappeared within a matter of seconds to mimic a more dynamic daily-life situation (Figure 1). We observed that humans capitalize on the reappearance of maintained items to boost another non-reappearing representation held in memory. Our findings are reminiscent of spatial and visual change-detection tasks that manipulated the similarity between the study and probe displays (Hollingworth, 2006, 2007; Jiang et al., 2000; Udale et al., 2018). Jiang et al. (2000) found that performance was best when probe displays were most similar to memorized locations or colors. These results seem to dovetail with the current findings from Experiments 2-4, as sensory input matching (visual) working memory content boosted performance. We extend these findings by presenting cues well before participants are able to respond, tapping into an internal prioritization process that prepares for upcoming behavior instead of the retrieval process itself. Together, this shows that the external world affects VWM operations beyond encoding, revealing that the environment is used to flexibly update the priority of memory content.

We repeatedly found that especially reappearing items fully matching memory facilitated VWM prioritization effectively (compared with items only matching in orientation or location). The sensory recruitment model of VWM may provide an explanation for this observation. The sensory recruitment account proposes that the perception and maintenance of sensory information recruits the same low-level sensory areas (D’Esposito, 2007; Gayet et al., 2018; Scimeca et al., 2018). Indeed, early visual areas are involved in perception as well as VWM (Harrison & Tong, 2009; Serences et al., 2009), and neural and pupil responses to visual input matching VWM content are enhanced (Gayet et al., 2017; Olmos-Solis et al., 2018; Wilschut & Mathôt, 2022) (also see Karabay et al., 2024). Maintaining material in VWM may pre-activate neuronal populations involved in processing specific visual features, leading to an enhanced neural response to matching input (Gayet et al., 2017). In the context of the current data, memorized orientations could have pre-activated orientation and spatially sensitive neurons during maintenance. Whenever items completely matching these pre-activated neuronal patterns reappear, they may be processed more efficiently compared with reappearing items that do not (fully) match these pre-activated patterns. In turn, enhanced processing of the reappearing material may help guide which remaining item should be prioritized. The sensory recruitment hypothesis also captures the results from Experiment 4 as interference caused by presenting novel orientations accounts for the observed accuracy benefits in the combination compared with the location conditions (also see Bays et al., 2024; Oberauer, 2002). Due to overlapping underlying neural areas, the presentation of novel non-memory matching grating may have cause interference to the memorized orientations, and this finding dovetails with work on distraction in VWM (e.g. Lorenc et al., 2021; Lout et al., 2023; van Moorselaar et al., 2015; Vogel et al., 2006; Wilschut & Mathôt, 2022). Instead when reappearing gratings matched VWM contents, it did not cause interference and instead boosted accuracy in a similar way to a spatial retro cue. Together, this indicates that the confluence of internal and external signals enable one to prioritize the most relevant information in the environment and memory.

Besides pre-activation through maintenance, another attentional process may have contributed to the boosted prioritization when fully memory-matching items reappeared. In van Ede et al. (2020), subtle gaze biases revealed that visual input during maintenance containing features (e.g., color or orientation) of items held in VWM can automatically ‘capture’ internal attention (see Ester & Nouri, 2023; Theeuwes, 1994) - even when these retro cues were uninformative. Such internal attentional capture possibly also occurred in our task, facilitating the use of reappearing items which led to enhanced prioritization. However, one could also argue the opposite for both the internal attentional capture and pre-activation accounts: If the memory-matching reappearing items are processed more deeply or automatically captured attention, this could distract from the most relevant VWM content that should be prioritized. We consider the latter possibility unlikely since reappearing items fully matching memory instead guided the prioritization process most effectively in our experiments. The reappearing items were temporarily task-relevant, and became obsolete only after extracting the relevant information. Thus, processing the reappearing items more efficiently aids the extraction of useful information to infer which item should be reported, leading to stronger prioritization of VWM content. Together, through the pre-activation of shared neural substrates and/or internal attentional capture, we posit that processing the reappearing items more deeply and rapidly is beneficial for performance.

Another possible mechanism at play in the current experiments are context-based effects. In line with recent proposals (e.g. Bays et al., 2024; Brady & Alvarez, 2011), it is possible that the reappearing items reactivated the ensemble of all memorized items. Such reactivation may have helped participants to “fill in” the to-be-probed item. One argument challenging this interpretation is that performance in the retro cue and reappear 2 was similar in Experiment 1. If context-based effects played a big role in the current results, one would perhaps expect participants to show more accurate and/or faster responses in the reappear 2 compared with the retro-cue condition. Nevertheless, this intriguing mechanism possibly underlying our effects cannot be ruled out given the current data, and is an interesting avenue for future investigations.

Let us briefly revisit the aforementioned example of crossing the street (Figure 1): our results indicate that seeing the red and the black car moving from left to right, facilitates the prioritization of the memorized black car. As van Ede and Nobre (2023) argue, many retro cue experiments may not reflect how memorized material is prioritized in natural settings, as one seldom receives explicit spatial or feature-based cues indicating what VWM content is most relevant. One way that memories may be prioritized more naturally is based on temporal expectations (Heuer & Rolfs, 2023; Nobre & van Ede, 2023; van Ede et al., 2017). Being able to predict *when* to act upon a specific representation allows to prioritize that representation which enhances its fidelity and protects it from interference (Gresch et al., 2022; Gresch et al., 2021; van Ede et al., 2017). In addition to temporal expectations, we here introduce another way how VWM prioritization may manifest itself in daily life: humans capitalize on the dynamic nature of sensory input to prioritize VWM content. However, the underlying internal prioritization mechanism do not necessarily differ between retro-cues, temporal expectations or the currently employed cues. It is indeed possible that the above examples are different ways to inform the same internal prioritization process to facilitate adaptive behavior.

Our findings open a number of intriguing questions. Although the underlying internal prioritization mechanism may very well be the same, investigating potential similarities and discrepancies between the here newly introduced reappearing-item cues and more traditional retro cues may shed light on how VWM content is prioritized in more natural situations. For example, what is the fate of the reappearing maintained items? It is possible that participants either selectively forget the reappearing items, or prioritized remaining material without dropping VWM content (Souza et al., 2014; M. Williams et al., 2013; M. Williams & Woodman, 2012). The current data cannot fully dissociate between these options because cues indicated to-be-reproduced items with full validity. However, the reappear-one condition from Experiment 1 provides a clue into this issue: If participants selectively dropped reappearing items, one would expect a benefit in this condition compared with the control condition. In contrast, although responses became faster, no clear benefit was found on accuracy as absolute errors were not significantly affected (and numerically even increased). Although we cannot make any claims regarding this point with certitude, this finding may indicate that instead of directed forgetting, participants used the reappearing items to internally attend another non-reappearing item held in memory, which some have hypothesized to be limited to a single representation (e.g. Olivers et al., 2011; van Moorselaar et al., 2014) (also see Souza & Oberauer, 2016) - although note this idea has been challenged (Beck et al., 2012; J. R. Williams et al., 2022) and the number of VWM representations guiding attention may be task-dependent (Frătescu et al., 2019; Wang et al., 2024). Future work may tackle whether forgetting plays a role by either implementing invalidly cued trials, as well as by independently manipulating set size and the number of reappearing items. Moreover, it is possible that reappearing items may retain relatively strong memory traces when compared with uncued items in designs with traditional retro cues because the items physically reappear, which may refresh the memory traces. These hypothesized stronger memory traces may manifest themselves in more accurate and faster responses when probing uncued items (i.e. less costs of cueing), having relatively strong biasing effects on reports of cued items, or even improving long-term memory representations. Additionally, it would be interesting to investigate whether the timecourse of internal prioritization would differ between more traditional retro-cues and reappearing-item cues. If participants would use higher level reasoning (e.g. deduction) when extracting the relevant information from the reappearing items, the ultimate prioritization process may occur relatively late when compared with selecting items from VWM based on their features or locations. Other open questions concern how reappearing items are used when their properties change across trials (i.e. mixed design) and how other types of information that reappearing items are able to convey, such as temporal order, may be used to guide VWM prioritization (as in Figure 1; also see Heuer & Rolfs, 2023).

Here, we investigated if, and how, reappearing task-irrelevant sensory input facilitates VWM prioritization. By demonstrating that humans capitalize on the dynamic nature of the world to prioritize VWM content, we advance our understanding of how memory prioritization may occur in more natural settings. Together, we propose that the relationship between VWM and sensory processing is bidirectional: VWM drives the selection of sensory input, which in turn guides the prioritization of the most relevant memory representations.

## Declarations

## Ethics approval

The experimental procedure was approved by the ethical review board of Utrecht University’s Faculty of Social Sciences (21-0297).

## Consent to participate

All participants provided written informed consent. Participants volunteered to take part in the experiment and were informed that they could abort the experiment whenever they wanted, and that they would be compensated for their time regardless of finishing or aborting the experiment.

## Consent for publication

All participants provided consent for publishing results based on their data as well as for the sharing of their data. Shared data are anonymous and personal details are omitted.

## Availability of Data and Materials

Analyses scripts and data are available via the Open Science Framework: https://osf.io/qzvkc/.

## Code Availability

Analyses scripts and data are available via the Open Science Framework: https://osf.io/qzvkc/.

## Acknowledgements

We thank Daan Gerits for his help with data collection for Experiments 2, 3 and 4, thank Surya Gayet for his help with the design of Experiment 3, and thank Daryl Fougnie for valuable discussions.

## Footnotes

1 We used the default settings, which include having a maximum number of 200 iterations, and estimates are obtained using the Broyden-Fletcher-Goldfarb-Shanno optimizer.

2 Note that our task does not offer a fair comparison between orientation and location information. As there were only three different locations, it may have been easier to use this information to guide internal prioritization compared with orientation which had 180 different orientations throughout the task. Thus, although participants seemed to not be able to use the isolated orientation information to guide internal prioritization here, it may be possible that in other designs orientation information may be used more effectively.

3 Note that this effect was non-significant when excluding the first author’s data, *t* = 1.69, *p* = .09.

